# Neocortical inhibitory interneuron subtypes display distinct responses to synchrony and rate of inputs

**DOI:** 10.1101/671248

**Authors:** Matthew M. Tran, Luke Y. Prince, Dorian Gray, Lydia Saad, Helen Chasiotis, Jeehyun Kwag, Michael M. Kohl, Blake A. Richards

## Abstract

Populations of neurons in the neocortex can carry information with both the synchrony and the rate of their spikes. However, it is unknown whether distinct subtypes of neurons in the cortical microcircuit are more sensitive to information carried by synchrony versus rate. Here, we address this question using patterned optical stimulation in slices of barrel cortex from transgenic mouse lines labelling distinct interneuron populations: fast-spiking parvalbumin-positive (PV+) and somatostatin-positive (SST+) interneurons. We use optical stimulation of channelrhodopsin-2 (ChR2) expressing excitatory neurons in layer 2/3 in order to encode a random 1-bit signal in either the synchrony or rate of activity in presynaptic cells. We then examine the mutual information between this 1-bit signal and the voltage and spiking responses in PV+ and SST+ interneurons. Generally, we find that both interneuron types carry more information than GFP negative (GFP-) control cells. More specifically, we find that for a synchrony encoding, PV+ interneurons carry more information in the first 5 milliseconds, while both interneuron subtypes carry more information than negative controls in their later response. We also find that for a rate encoding, SST+ interneurons carry more information than either PV+ or negative controls after several milliseconds. These data demonstrate that inhibitory interneuron subtypes in the neocortex have distinct responses to information carried by synchrony versus rates of spiking.

## Introduction

One of the foundational concepts in neuroscience is that neurons encode information with their action potentials. The earliest demonstrations that action potentials carry information came from Lord Adrian (1926), who demonstrated that the rate of spikes in peripheral nerves is correlated with the force applied to a limb^1^. Decades of work following this demonstrated that the rate of spikes can carry information about almost any aspect of the environment or an animal’s behaviour, including sensory stimuli^2^, spatial location^3^, action selection^4^, etc. However, the rate-of-fire of a neuron is not the only aspect of a spike train that can carry information^5^. It has also been demonstrated that the specific timing of action potentials can correlate with salient variables, including sensory stimuli^6,7^, spatial location^8^, and action selection^9,10^. There are a variety of potential coding schemes using spike times^5^, but basic principles of postsynaptic spatiotemporal integration tell us that the synchrony of incoming inputs can be as important as the rate of incoming inputs to a neuron^11^. Therefore, the brain is likely to encode information using both the synchrony and rate of spikes^12–14^.

One interesting aspect of information encoding with both synchrony and rate of spikes is that different cells with different biophysical properties will respond to each signal differently^15,16^. For example, when we consider linear integration, a cell with a short membrane time constant will be more sensitive to information carried by synchronous inputs, whereas a cell with a longer time constant would be more sensitive to the rate of inputs. These issues are particularly salient when we consider the diversity of biophysical properties found in neocortical inhibitory interneurons^17–19^. Different types of inhibitory interneurons possess intrinsic membrane properties, morphologies, firing patterns, and presynaptic inputs^17,19^. For example, parvalbumin-positive (PV+) inhibitory interneurons are known to have very short membrane time constants, fast-spiking behaviour, and short-term depressing presynaptic inputs^20^. In contrast, somatostatin-positive (SST+) inhibitory interneurons possess adapting firing patterns and short-term facilitating presynaptic inputs^21^. These distinct biophysical properties of interneurons are likely relevant to information encoding in the brain^16,22,23^.

Given that different interneuron subtypes, such as PV+ and SST+ cells, display distinct biophysical properties, it is possible that each subtype is more or less sensitive to information conveyed by presynaptic synchrony or rate. However, although interneuron subtypes, particularly PV+ and SST+, have been studied extensively over recent decades, it is unknown whether there are functional specializations in the integration of information carried by spike synchrony or spike rate.

Here, we explored this question using a combination of transgenics, *ex vivo* whole-cell patch clamping, and patterned optogenetic illumination. Specifically, we examined the responses of PV+ and SST+ inhibitory interneurons in the barrel cortex of mice when presynaptic excitatory inputs were driven optogenetically. Using a digital multimirror device, we encoded a random 1-bit signal by controlling either the synchrony or rates of optical activation of presynaptic inputs. We then examined the amount of information encoded by PV+, SST+ and pyramidal neurons about this 1-bit signal. We found that these two classes of cell were differentially sensitive to the synchrony and rate of optical activations of presynaptic inputs. Specifically, we observed that both interneuron types carried more information in their spiking responses than pyramidal neurons but in different ways. PV+ interneurons carried more information about the 1-bit signal in their early (<5 ms) responses to synchronous activation. Both interneuron types carried more information in response to synchronous activation after more time (> 5ms). When we examined the responses to rate encoding, SST+ interneurons carried more information about the 1-bit signal than either PV+ or pyramidal neurons in their later responses. These data confirm that different types of inhibitory interneurons can integrate information carried by synchrony or rate of inputs in different ways. It suggests that the inhibition received by pyramidal neurons in the neocortex may be selectively driven by the synchrony and the rate of action potentials.

## Results

### Transgenic targeting of PV+ and SST+ interneurons

We focused our investigation on layer 2/3 the somatosensory cortex, as this is a region of the neocortex where synchrony and rate codes have been extensively explored. In order to target different populations of interneurons within L2/3 of the barrel cortex, we used GAD67-GFP and Gin-GFP transgenic mice which have been reported to selectively express GFP in PV+ and SST+ interneurons, respectively^24,25^. To confirm this selective expression of GFP, we characterized GFP cells in these two transgenic lines using immunohistochemical and electrophysiological properties.

Immunohistochemistry staining for PV protein in the barrel cortex of GAD-67-GFP animals showed a higher probability of GFP+ cells being PV+ (P(PV+|GFP+), 59.4% ± 7.5%, *n* = 7 animals) and vice versa (P(GFP+|PV+), 33.5% ± 3.1%, *n* = 7) when compared to staining for SST peptide (P(GFP+|SST+), 5.0% ± 1.4%, *n* = 6; P(SST+|GFP+), 3.0% ± 0.6%, *n* = 6; **Fig. 1A** *top row*). In Gin-GFP animals, the opposite was seen. In particular, when staining for SST, Gin-GFP animals showed a higher probability of GFP+ cells being SST+ (P(SST+|GFP+), 94.9% ± 7.0%, *n* = 5) and vice versa (P(GFP+|SST+), 74.3% ± 4.2%), than when compared to PV protein staining (P(PV+|GFP+), 0.2% ± 0.2% *n* = 6, P(GFP+|PV+), 0.1% ± 0.1, *n* = 6; **Fig. 1A**, *bottom row*). These results suggest a reasonable level of both genotypic specificity and efficiency and is in line with previous work which suggests that GAD67-GFP and Gin-GFP animals express GFP in PV+ and SST+ cells, respectively^24–27^.

**Figure 1.**
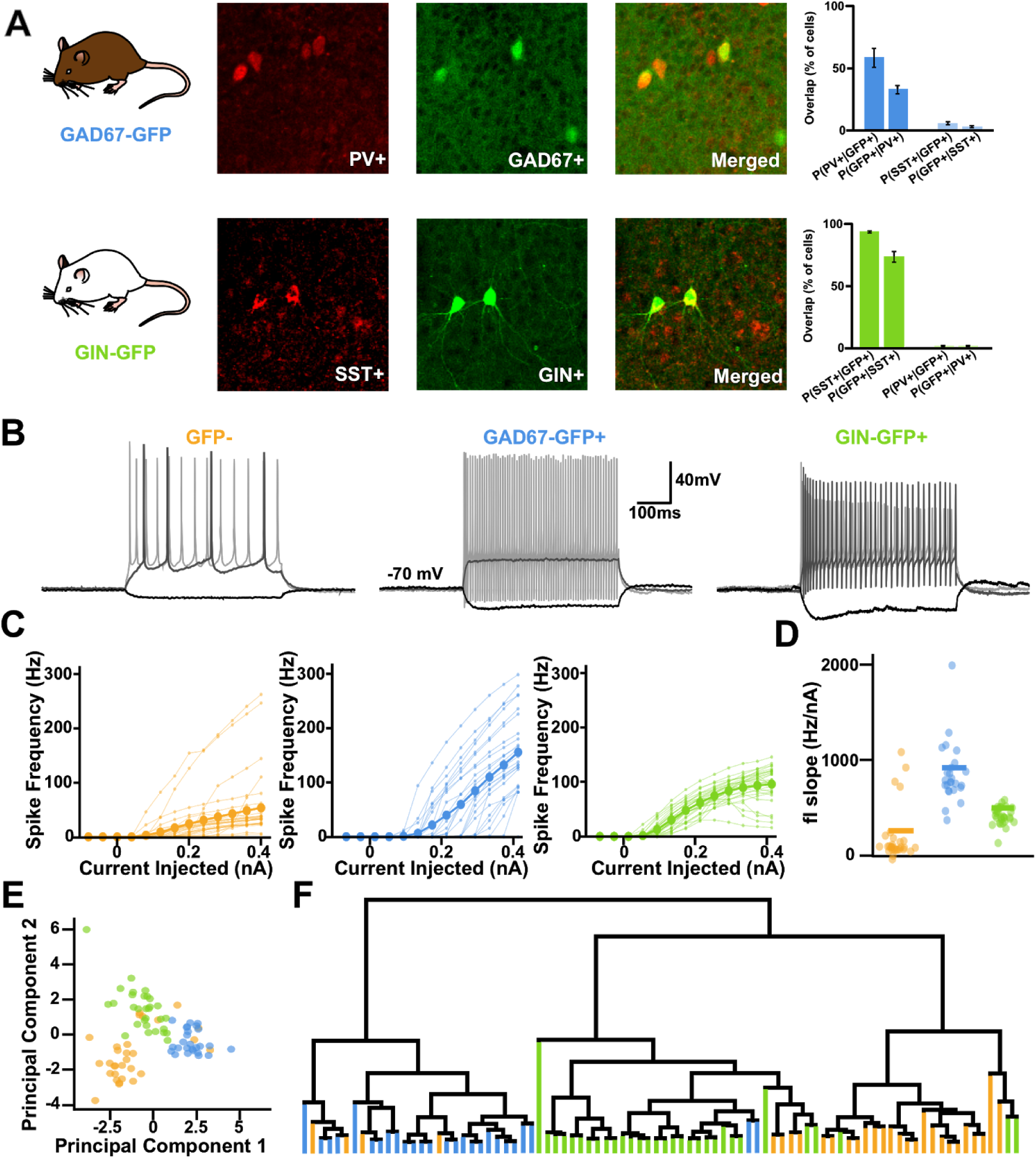
Gad67-GFP and Gin-GFP mice target distinct populations of neurons, likely fast-spiking PV+ and SST+ interneurons, respectively. A) Sample immunohistochemistry images from both transgenic lines. B) Sample electrophysiological traces from each cell type to current injections −80 pA, +160 pA, +400 pA (darkest to lightest). C) Frequency vs. current injection (fi) curves for each cell type. D) Mean slope of the fi curves for each cell type. Larger circles represent mean values. E) 2 Component PCA of each cell type’s electrophysiological characteristics. F) Dendrogram clustering.

Whole-cell patch clamp current recordings from both GFP- and GFP+ cells in layer 2/3 of the barrel cortex were also used to analyze each cell’s electrophysiological properties. In the GAD67-GFP transgenic line, nearly all of the GFP+ cells (N=26) exhibited spiking behaviour typical of cortical PV+ cells: they were often fast-spiking and reached average peak frequencies of roughly 150 Hz or more in response to positive current clamp injections (Fig 1B,C,D). GFP+ cells in the Gin-GFP transgenic line (*n* = 30) showed different spiking characteristics to the GAD67-GFP+ cells as their spiking tended to accommodate in response to increasing current injections and often peaked at around 100 Hz (**Fig. 1B-D**), a hallmark firing pattern for almost 90% of SST+ cells^26^.

To compare with these cell populations, we also recorded from GFP-cells in each transgenic line (*n* = 30). Usually, these GFP-cells exhibited regular firing patterns typical of layer 2/3 pyramidal cells (**Fig. 1B-D**). Sometimes, they exhibited fast-spiking behaviour (4 cells), which is to be expected from a random sampling of cells in the circuit. These four cells are shown here, but were excluded from later analyses. A principal components analysis (PCA) of each cell’s electrophysiological characteristics (e.g. adaptation ratio, sag amplitude, and membrane tau; see *Materials and Methods*) revealed three distinct clusters in the first two principal components (**Fig. 1E**). When these components were used to map the recorded cells onto a dendrogram, three distinct clusters that were largely consistent with selective labelling of distinct subtypes were revealed (**Fig. 1F**).

Altogether, these results confirm that GAD67-GFP and Gin-GFP transgenic mice express GFP in distinct cell types, likely corresponding to PV+ and SST+ interneurons and that most GFP-cells are not from these subclasses of interneurons.

### Encoding a random 1-bit signal using patterned optical stimulation

Central to our experimental objective is the ability to encode information via the synchrony or rate of presynaptic inputs to a neuron. To do this, we adopted a patterned optical stimulation approach. We infected the barrel cortex of 5-7 week old mice with an adeno-associated virus carrying channelrhodopsin-2 (ChR2) and the mCherry reporter, under the Ca^2+^/calmodulin-dependent protein kinase II promoter (rAAV1-CamKii-hChR2(h134r)-mCherry). This led to expression in pyramidal neurons of layer 2/3, and we did not observe any ChR2 expression in GFP+ neurons (**Fig. 2A**). After sufficient time for expression (2-3 weeks), we then prepared *ex vivo* slices of barrel cortex and used a digital multimirror device to illuminate layer 2/3 with spatially controlled patterns of light (**Fig. 2B**; see *Materials and Methods*). Whole-cell recordings from infected neurons demonstrated that we could use spatially restricted discs of illumination (470 nm, 15 μm diameter, ∼14 mW/mm^2^) to reliably induce spiking in targeted pyramidal neurons (**Fig. 2C**). We also examined the responses of neurons at different distances from the disc of illumination (**Fig. S1A**). We found that due to the limitations of 1-photon excitation, neurons up to 50 μm away from a disc of illumination could also spike (**Fig. S1B-D**). Nonetheless, targeted neurons spiked earlier and more reliably than non-targeted neurons, and neurons farther than 50 μm rarely spiked (**Fig. S1B-D**). This demonstrated that we could use the digital multimirror device to selectively activate particular regions of interest (ROIs) in the slice.

**Figure 2.**
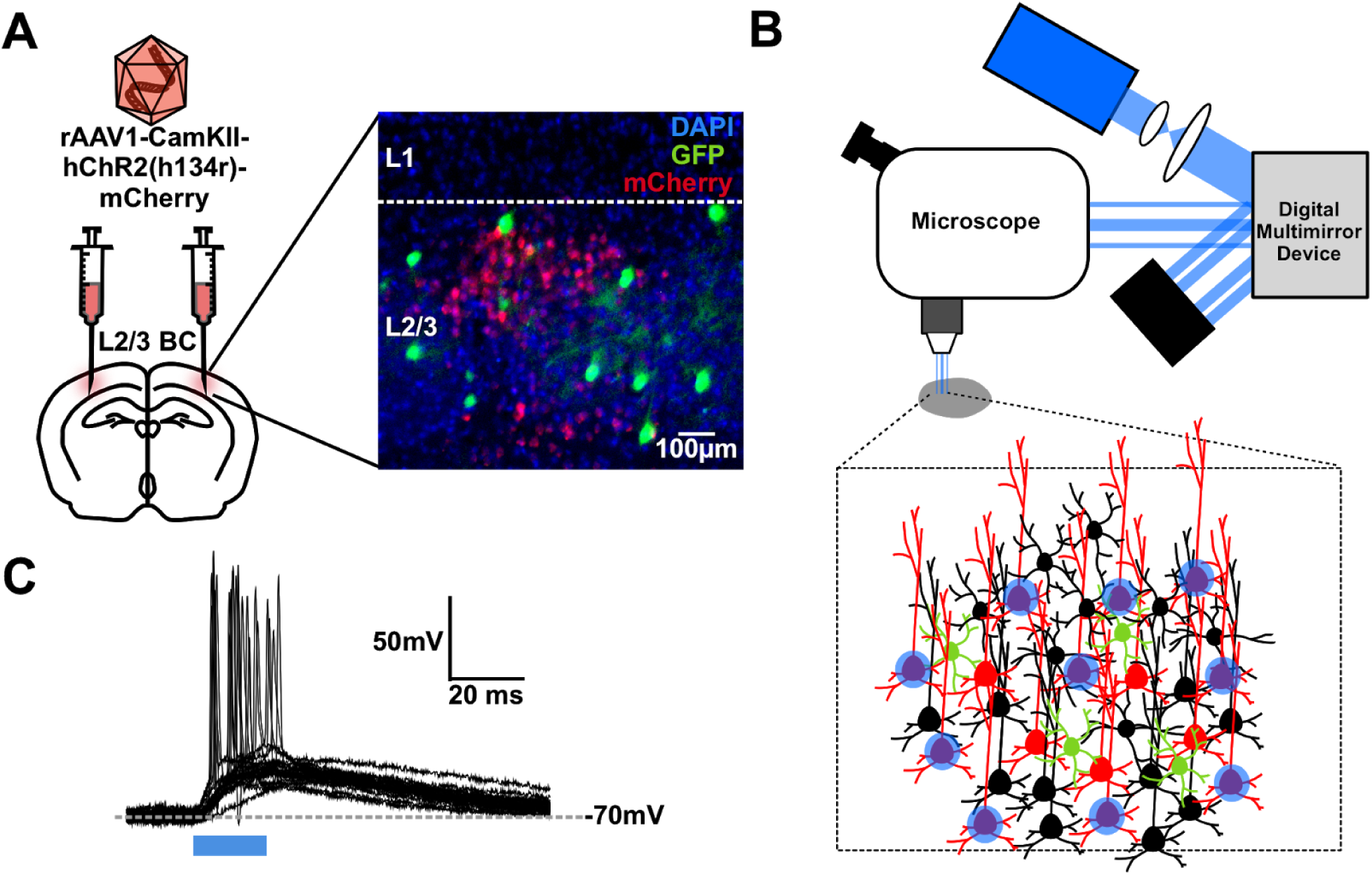
Patterned optical stimulation of ChR2+ Layer 2/3 Pyramidal Neurons. A) Viral transfection of Channelrhodopsin-2 into L2/3 Pyramidal Neurons. B) Illustration of patterned optical stimulation protocol. C) Sample responses from ChR2+ layer 2/3 Pyramidal neurons (n=15) to a 15 μm spot placed directly on top of the recorded neuron.

Next, we performed whole-cell patch clamp recordings of GFP+ (and some GFP−) neurons in the slices. We used our patterned optical illumination approach to encode a 1-bit random signal (i.e. a signal with two states, 0 or 1) in the activity of 10 ROIs containing ChR2 expressing neurons that were presynaptic to the patched neuron (**Fig. 3A-B**). We encoded this signal using either the synchrony or rate of optical activation of the 10 presynaptic ROIs. To restrict our data to 10 ROIs with monosynaptic inputs to the patched neuron, we examined the postsynaptic responses in the first 5 ms of optical activation (**Fig. 3B**, *insets*). We only included recordings for subsequent analyses if the 10 ROIs elicited monosynaptic-driven depolarization, i.e. depolarization commencing 0.5-2 ms after stimulation onset, in at least 90% of the optical stimulations (**Fig. 3C**; see *Materials and Methods*).

**Figure 3.**
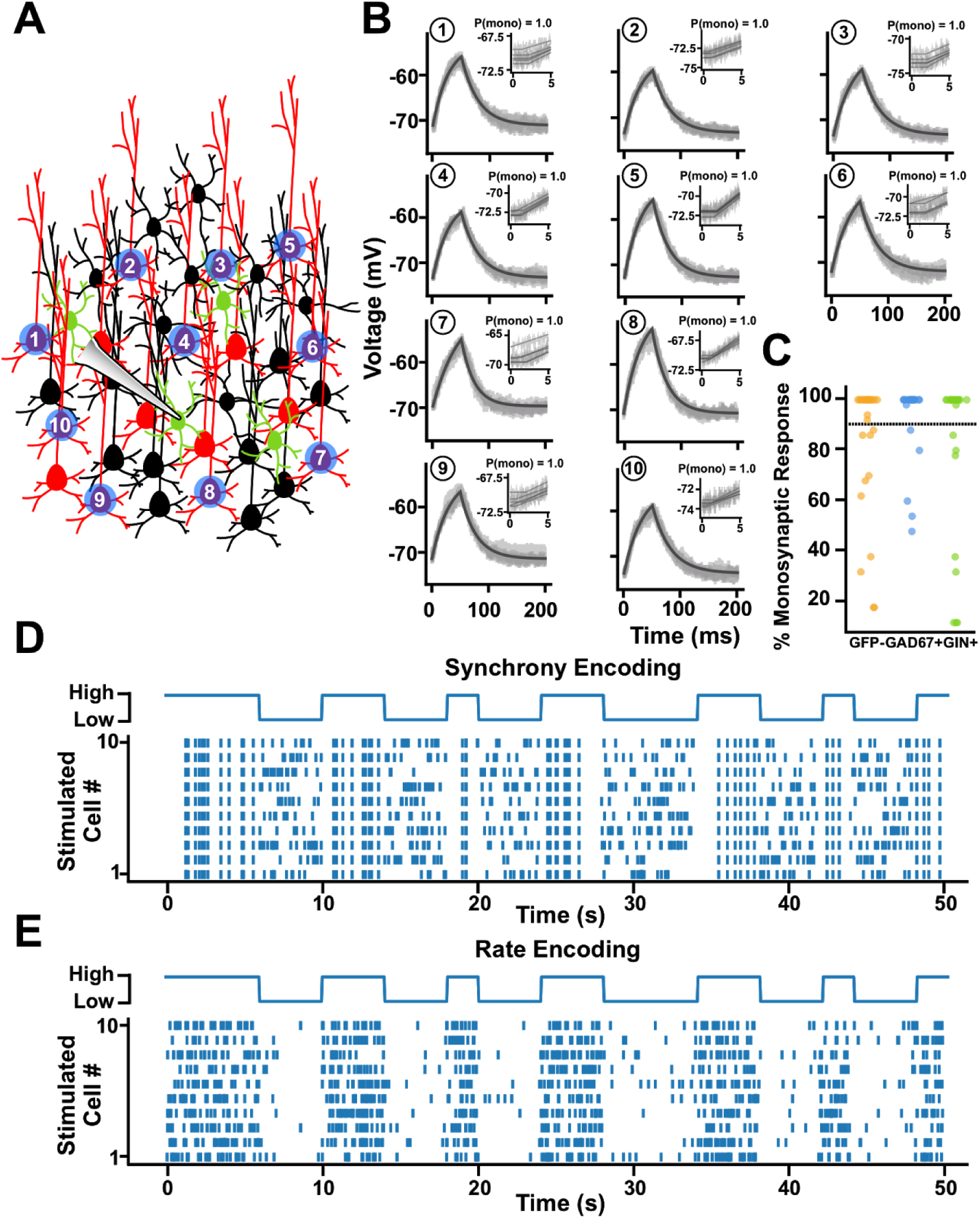
Artificially encoding a random 1-bit signal using the synchrony or rate of optical inputs to ChR2+ Pyramidal Neurons in Layer 2/3. A) Experimental protocol. B) Example postsynaptic traces from a sample cell showing 10 presynaptic cells which passed the monosynaptic reliability test. C) Distribution of input cells and their monosynaptic reliability. D) Sample raster plots illustrating how the 1-bit signal was encoded using synchrony of optical inputs. E) Sample raster plots illustrating how the 1-bit signal was encoded using the rate of optical inputs.

To encode the 1-bit signal using the synchrony of activation, we created an optical activation pattern where each ROI was activated at a constant rate of 2.7 Hz, but with low synchrony for the 0 state, and high synchrony for the 1 state (**Fig. 3D**). Specifically, during the low synchrony state, the activation times of the 10 ROIs were sampled from 10 independent Poisson processes, whereas during high synchrony states the activation times were sampled from a single Poisson process. As a result, when the 1-bit signal was in the 0 state, the 10 ROIs were activated at independent times, whereas when the 1-bit signal was in the 1 state, the 10 ROIs were activated synchronously. But, importantly, the rate of activation of the ROIs was identical in the two states.

To encode the 1-bit signal using the rate of activation, we used a low rate for the 0 state and a high rate for the 1 state (**Fig. 3E**). Specifically, we always sampled the activation times of the ROIs from 10 independent Poisson processes, but for the 0 state we sampled with a 0.5 Hz rate, and for the 1 state we sampled with a 5 Hz rate. This meant that the average rates were approximately the same as for the synchrony encoding (2.7 Hz), but the rates changed depending on the state of the 1-bit signal. Interestingly, we found that, in comparison to whole-field illumination protocols, these patterned optical illumination protocols produced qualitatively more natural responses in the recorded neurons, which was confirmed by analyzing the spectral densities of the whole-cell recordings (**Fig. S2**). Therefore, using our synchrony and rate encoding protocols, we could investigate the extent to which different subtypes of neurons in layer 2/3 barrel cortex are sensitive to information encoded with the synchrony or rate of presynaptic inputs.

### Responses to synchrony encoding differ between neuron subtypes

We first examined the responses of GAD67-GFP+ cells (likely PV+ interneurons), Gin-GFP+ cells (likely SST+ interneurons), and non-fast spiking GFP-, non-fluorescent (NF) cells (likely pyramidal neurons) to the synchrony encoding protocol (NF: *n* = 17, PV+: *n* = 21, SST+: *n* = 22). In general, all three types of neurons exhibited reliable responses to the optical stimulation patterns (**Fig. 4A**). Histograms of both the mean membrane potential and the firing frequency (i.e. spike counts) for each cell type over the course of the 50 ms window illustrated different responses during the 0 and 1 states of the random signal. As expected, we found that both average membrane potential and spiking frequency were variable and correlated with the number of presynaptic ROIs that were activated, such that during the 0 state of low synchrony the spike counts and membrane potential were highly variable, but during the 1 state of high synchrony, the spike counts and membrane potential were less noisy (both were high when all the ROIs were activated, and low when none of the ROIs were activated). More specifically, in the 0 state, the average membrane potential of all three cell types were generally more depolarized and variable (**Fig. 4B, light histograms**; NF = −58.98 mV ± 4.54, PV+ = −59.51 mV ± 4.36, SST+ = −54.96 mV ± 4.92) when compared to neurons in the 1 state (**Fig. 4B, dark histograms**; NF = −65.47 mV ± 2.15, PV+ = −66.37 mV ± 2.24, SST+ = −64.35 mV ± 2.89). Similarly, we found that during the 0 state of low synchrony, the firing frequency across each cell type was generally higher and extremely variable (**Fig. 4C, light histograms**; NF = 3.29 Hz ± 2.80, PV+ = 19.58 Hz ± 17.87, SST+ = 15.01 Hz ± 8.31), compared to spiking frequencies during the 1 state of high synchrony, which were more consistent (**Fig. 4C, dark histograms**; NF = 2.59 Hz ± 1.81, PV+ = 9.84 Hz ± 5.67, SST+ = 8.47 Hz ± 1.83).

**Figure 4.**
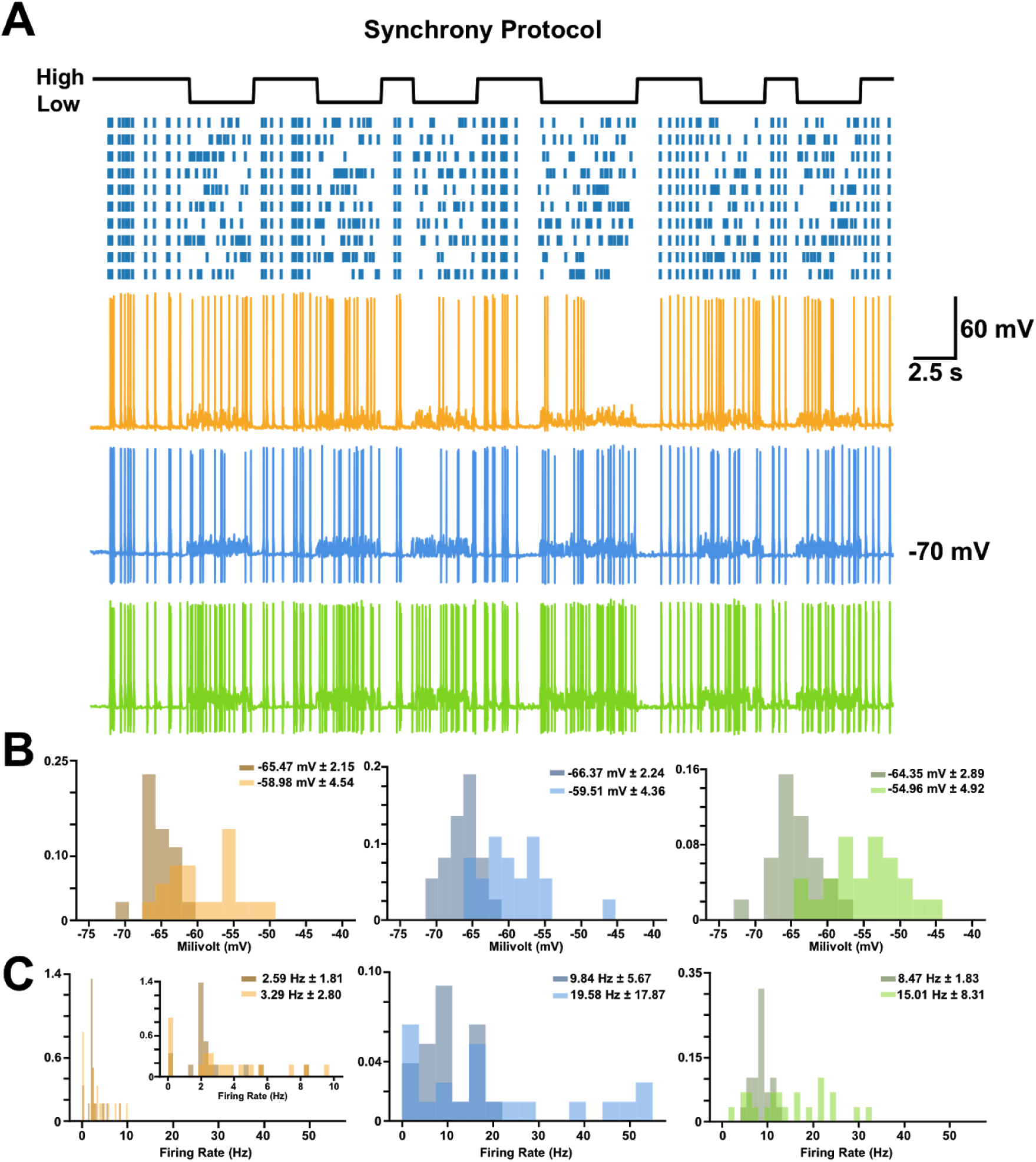
High and low states within the synchrony encodings produced different neuronal responses. A) Sample traces of each cell type to the synchrony encoding. B) Probability density functions of the membrane potential of each cell type during high (darker histogram) and low (lighter histogram) states within the synchrony encodings (numbers shown represent mean membrane potential in each state ± s.d.). C) Probability density functions of the firing frequency of each cell type during high and low states within the synchrony encodings (numbers shown represent mean firing frequency in each state ± s.d.).

To determine the extent to which each subtype of neuron was sensitive to information encoded in the synchrony of presynaptic inputs, we used information theoretic tools. Specifically, we examined the mutual information between the 1-bit signal and different aspects of the patched cells’ activity. Because our optical protocol did not include any inhibition, the activation of the 10 ROIs induced prolonged polysynaptic activity in the tissue (**Fig. S3**). As such, in order to estimate potential differences in monosynaptic versus polysynaptic responses, we divided our analyses into early responses found in the first 5 ms of optical activation (which should be largely monosynaptic) and later responses (which should be largely polysynaptic).

When we analyzed whether the average membrane potential of each cell type conveyed information about the random signal, we found that each neuron type was roughly equal in the amount of mutual information between the 1-bit signal and their mean voltage, for both the monosynaptic (**Fig. 5A**; NF = 0.3928 ± 0.1294, PV+ = 0.4206 ± 0.1075, SST+ = 0.4211 ± 0.0827) and polysynaptic windows (**Fig. 5B**; NF = 0.4562 ± 0.1353, PV+ = 0.4917 ± 0.0796, SST+ = 0.5079 ± 0.0681). This data suggests that although there may be differences in the responses of each neuron type to the synchrony encoded information, the amount of information they carry in their average membrane potential is equivalent.

**Figure 5.**
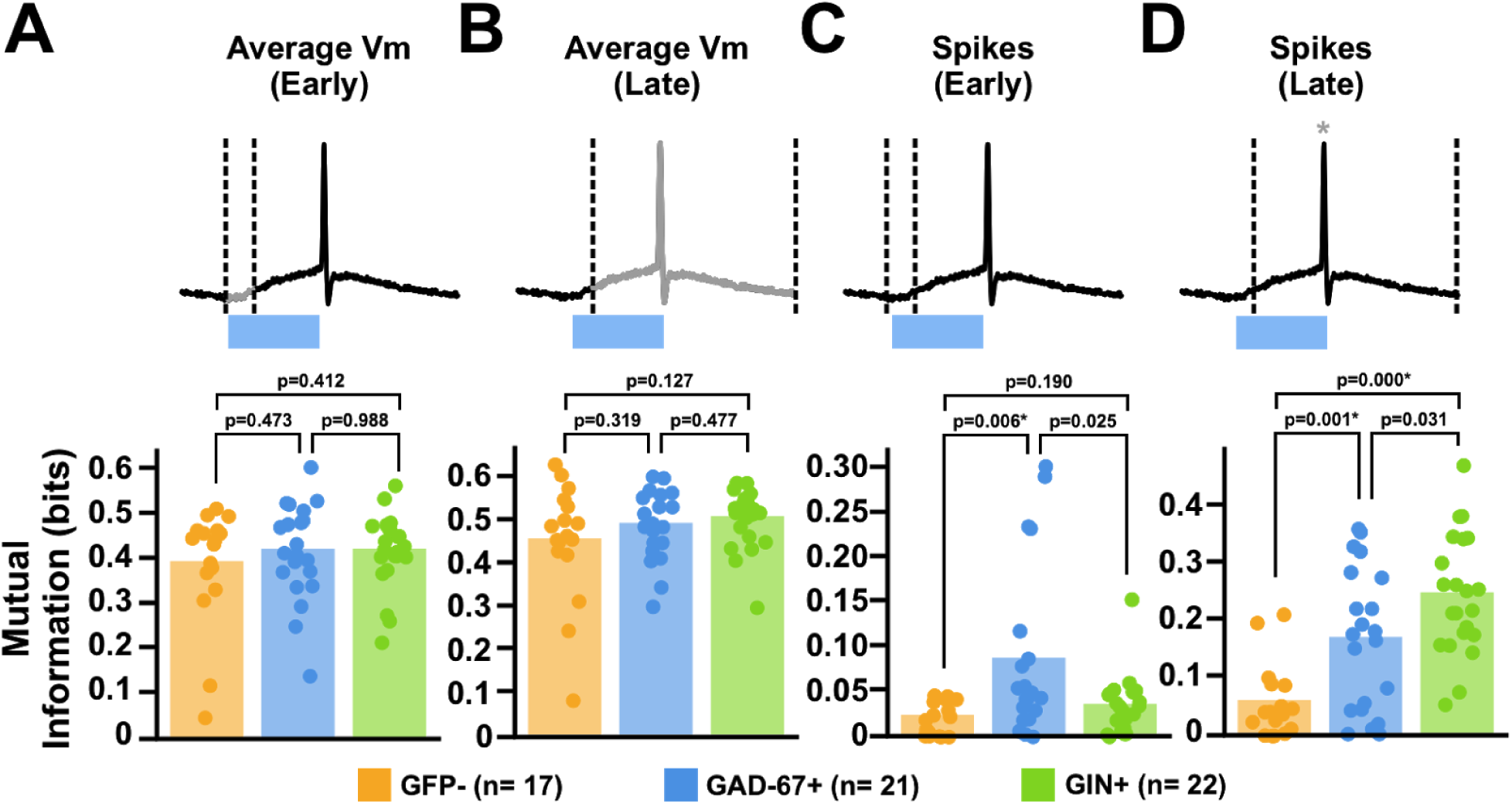
Different subtypes of interneurons carry different amounts of information in response to our synchrony encoding. A) Mutual information analysis of the average membrane potential of each cell type to our 1-bit signal within early (0-5ms) response. Dotted lines indicate window of analysis, with greyed areas indicating what part of the response was analyzed. B) Same analysis as C) but restricting the analysis to only the later (5-50ms) responses. C) Mutual information analysis of the spike counts of each cell type to our 1-bit signal within early (0-5ms) response. D) Same analysis as C) but restricting the analysis to only the later (5-50ms) responses.

Next, we examined the spiking responses of the patched neurons. Again, we split our analyses into approximately monosynaptic and polysynaptic time windows. Interestingly, unlike the mean voltage responses, we observed clear differences between neuron types in the mutual information between the 1-bit signal and the spike counts. In the monosynaptic window, GAD67-GFP+ cells showed the highest amount of mutual information with the random 1-bit signal (**Fig. 5C**; NF = 0.0218 ± 0.0187, PV+ = 0.0861 ± 0.0954, SST+ = 0.0338 ± 0.0332). These data suggest that when we consider spiking behaviour, PV+ interneurons rapidly convey more information than either SST+ interneurons or pyramidal neurons about signals encoded with synchronous monosynaptic inputs.

Meanwhile, for the polysynaptic time window, the GAD67-GFP+ and Gin-GFP+ cells showed equal levels of mutual information with the 1-bit signal, both of which were higher than the information contained in the spike counts of the GFP-neurons (**Fig. 5D**; NF = 0.0580 ± 0.0641, PV+ = 0.1688 ± 0.1242, SST+ = 0.2472 ± 0.1049 NF = 0.0580 ± 0.0641, PV+ = 0.1679 ± 0.1254, SST+ = 0.2472 ± 0.1049). This suggests that both interneuron subtypes may be more sensitive to signals encoded with synchronous inputs than pyramidal neurons. However, it should be noted that in the polysynaptic time window there may be differences in the rate of synaptic inputs thanks to the reverberation of activity in the tissue. This could include not only the 10 ROIs themselves but also neurons activated by the 10 ROIs. So, whether these differences reflect a different sensitivity to synchrony of inputs, or different sensitivity to inputs driven by synchronous inputs, is impossible to know. Nonetheless, taken together with the data from the monosynaptic time window, we can say that our results demonstrate that PV+ interneurons, SST+ interneurons, and pyramidal neurons carry different amounts of information about a 1-bit signal that has been encoded via optical activation of 10 ROIs in a synchronous vs. non-synchronous manner.

### Responses to a rate encoding differ between neuron subtypes

We then examined responses (NF: *n* = 17, PV+: *n* = 20, SST+: *n* = 22) to the same one-bit signal encoded via the rate of activation of the ROIs. Similar to responses seen in the synchrony optical encoding, each cell type showed different responses during the 0 or 1 state of the rate encoding, except that now the responses during the 0 state (low rate) were lower magnitude and less noisy than the responses during the 1 state (high rate) (**Fig. 6A**). More specifically, membrane potentials during the 0 state were often hyperpolarized and low variance (**Fig 6B, light histograms**; NF = −66.38 mV ± 2.15, PV+ = −67.38 mV ± 2.50, SST+ = −65.81 mV ± 2.34) whereas during the 1 state they were depolarized with high variance (**Fig. 6B, dark histograms**; NF = −54.71 mV ± 5.29, PV+ = −56.03 mV ± 6.67, SST+ = −49.06 mV ± 4.30). This pattern was also seen with the firing rates of each cell type in response to low rates of optical input (**Fig. 6C, light histograms**; NF = 0.90 Hz ± 0.74, PV+ = 4.61 Hz ± 4.79, SST+ = 4.68 Hz ± 2.56) versus high rates (**Fig 6C, dark histograms**; NF = 6.56 Hz ± 5.20, PV+ = 32.65 Hz ± 31.49, SST+ = 22.57 Hz ± 8.41).

**Figure 6.**
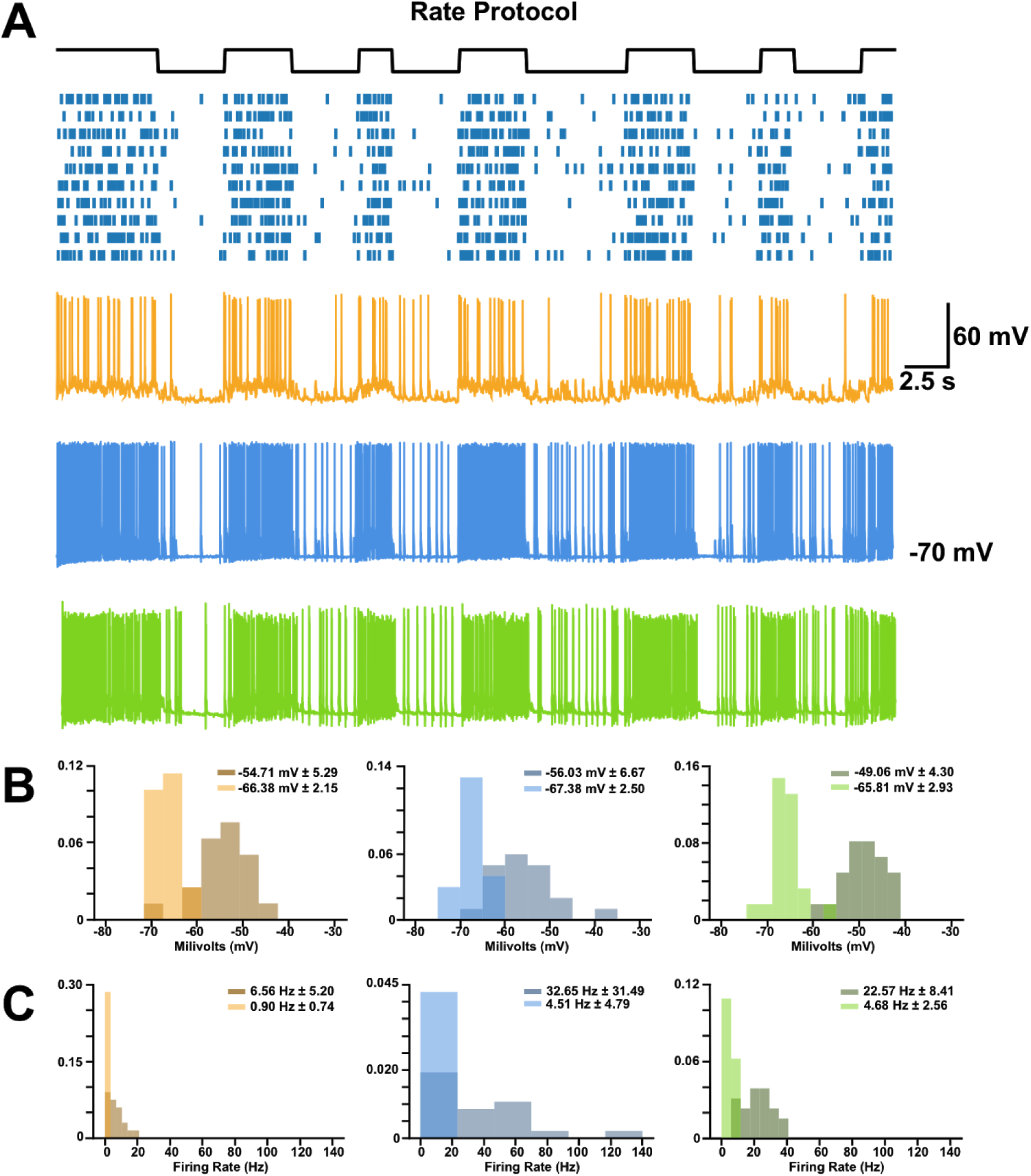
High and low states within the rate encodings produced different neuronal responses. A) Sample traces of each cell type to the rate encoding. B) Probability density functions of the membrane potential of each cell type during high (darker histogram) and low (lighter histogram) states within the rate encodings (numbers shown represent mean membrane potential in each state ± standard deviation). C) Probability density functions of the firing frequency of each cell type during high and low states within the rate encodings (numbers shown represent mean firing frequency in each state ± standard deviation).

Before conducting information theoretic analyses, we sought to ensure that any information about the signal encoded by the postsynaptic response was driven by the increase in the rate of optogenetic activation, rather than the unavoidable increase in synchronous stimulation of ROIs with increased rate (see **Fig. S4**). As such we conditioned our mutual information measure on the ROI activation count in each time bin (see *Materials and Methods*).

Conditional mutual information analysis revealed that PV+ interneurons tended to have lower conditional mutual information with the 1-bit signal than the other cell types in both the monosynaptic (**Fig. 7A**; NF = 0.1682 ± 0.0286, PV+ = 0.1161 ± 0.0519, SST+ = 0.1755 ± 0.0278) and polysynaptic (**Fig. 7B**; NF = 0.0802 ± 0.0238, PV+ = 0.0510 ± 0.0253, SST+ = 0.0782 ± 0.0214) windows. These data indicate that the membrane potential fluctuations of PV+ interneurons are less sensitive to rate encoded signals than SST+ interneurons and pyramidal cells.

**Figure 7.**
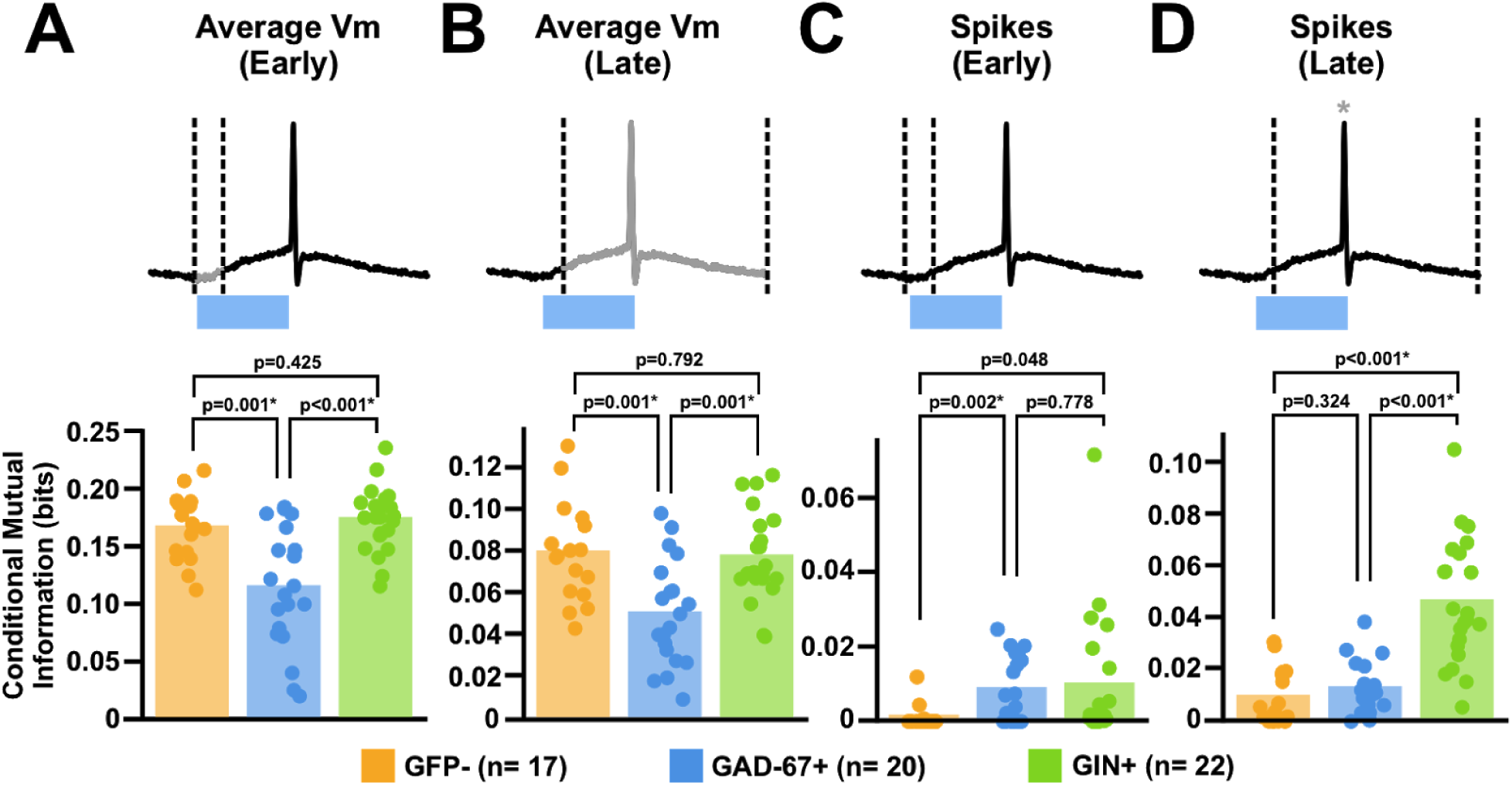
Different subtypes of interneurons carry different amounts of information in response to our rate encoding. A) Mutual information analysis of the average membrane potential of each cell type to our 1-bit signal within early (0-5ms) response. Dotted lines indicate window of analysis, with greyed areas indicating what part of the response was analyzed. B) Same analysis as A) but restricting the analysis to only the later (5-50ms) responses. C) Mutual information analysis of the spike counts of each cell type to our 1-bit signal within early (0-5ms) response. D) Same analysis as C) but restricting the analysis to only the later (5-50ms) responses.

We then examined the spiking responses of our recorded neurons, again splitting spike times into those occurring during the monosynaptic and postsynaptic windows. Spiking was more frequent in the 1 state than in the 0 state for all neurons, and spike counts tended to increase monotonically with the number of ROIs activated. The purpose of conditioning on ROI activation count is to determine whether there was a difference in spike counts accounted for only by the rate of inputs, and not the number of ROIs activated. We found that for spikes occurring during the monosynaptic window, there was little difference in conditional mutual information with the one-bit signal between PV+ and SST+ interneurons, although both tended to carry more information about the signal than NF cells (**Fig. 7C**; NF = 0.0010 ± 0.0030, PV+ = 0.0085 ± 0.0089, SST+ = 0.0097 ± 0.0172). This indicates that rapid responses to rate encoded pyramidal cell activity are similar between PV+ and SST+ interneurons.

Interestingly, for spikes occurring during the polysynaptic time window, SST+ interneurons carried more information about the signal than both the PV+ and NF cells, which carried similar amounts of information (**Fig. 7D**; NF = 0.0091 ± 0.0106, PV+ = 0.0126 ± 0.0102, SST+ = 0.0463 ± 0.0243). This indicates that SST+ interneurons can accumulate information about rate encoded signals over longer time windows than both PV+ and NF cells.

## Discussion

Using a digital multimirror device, we performed *ex vivo* experiments in slices of mouse barrel cortex to examine the sensitivity of different interneuron subtypes to information encoded with the synchrony or rate of presynaptic inputs. Using GAD67-GFP and Gin-GFP transgenic mice coupled with viral infection of pyramidal neurons with ChR2, we were able to examine the responses of layer 2/3 PV+ and SST+ interneurons (as well as NF, GFP-cells that were likely pyramidal neurons, **Fig. 1**). We examined their responses to a 1-bit random signal encoded with either the synchrony or the rate of optical ROI activation (**Fig. 2-3**). We found that there were indeed differences between cell types in the amount of information carried about the 1-bit signal. When the signal was encoded using the synchrony of ROI activation, all of the cell types carried similar amounts of information in their membrane potentials, but spiking responses showed differences (**Fig. 4-5**). PV+ interneurons carried more information than the other cell types during an early, likely monosynaptic time-window, while both PV+ and SST+ interneurons carried more information than NF cells in a later time-window. When the signal was encoded with the rate of ROI activation, we found that PV+ interneurons carried less information than either SST+ or NF cells in their membrane potential. For spiking responses, both PV+ interneurons and SST+ interneurons carried more information than NF cells in the early monosynaptic window, but in the later time-window, SST+ interneurons carried more information than either NF or PV+ cells (**Fig. 6-7**). Altogether, these results demonstrate that there are differences between neocortical cell types in their sensitivity to information encoded with the synchrony or rate of presynaptic inputs.

Our findings are broadly in-line with what is known about neuronal subtypes in the neocortex. First, we found that both interneuron types tended to carry more information than the NF cells in their spiking responses. This fits with a previous study which reported higher amounts of information about sensory stimuli in barrel cortex inhibitory interneurons compared to excitatory neurons^28^. (Though it should be noted that that study reported no significant difference in information content between interneurons and excitatory neurons in layer 2/3, which we have studied here.) Second, our finding that PV+ interneurons’ spikes rapidly convey information about a signal encoded with synchronous inputs, while SST+ interneurons gradually accumulate information about a signal encoded with different rates of inputs, fits with what is broadly known about the biophysics of these cell types. Specifically, the rapid membrane time-constants, short-term depressing synaptic inputs, and rapid spiking properties of PV+ interneurons^20^ fits with rapid transmission of information about synchronous inputs, while the adaptive spiking responses and short-term facilitating synaptic inputs of SST+ interneurons^26^ fits with long latency responses to high rate inputs.

Our data suggest that there are potential divisions of labour in information encoding in the neocortical microcircuit. Both PV+ and SST+ subtypes carried information about the1-bit signal regardless of the encoding format we used, but our results imply that, roughly, PV+ interneurons rapidly provide more information about synchronous inputs, while SST+ interneurons gradually accumulate information about the rate of inputs. Given this, the synchronous activation of excitatory neurons in the circuit^29,30^ may activate perisomatic inhibition more strongly, while recurrent or top-down signals that build-up over time may activate distal dendritic inhibition more strongly. This would also suggest that reports that synchrony and rate can carry distinct information about sensory stimuli^9,12^ may link certain aspects of sensory stimuli with certain forms of inhibition to pyramidal neurons. Future work should examine whether the information carried by PV+ and SST+ interneurons reflects signals communicated by the synchrony and rate of pyramidal cell activation, respectively.

It is important to note that our study was limited by a number of factors. First, we were performing our experiments *ex vivo*, and there are likely important differences in interneuron activity *in vivo* that relate to things like movement or neuromodulation^25^. Second, because we were using single-photon excitation with no optical inhibition, we could not guarantee that only a single neuron was being activated by illumination of the ROIs (**Fig. S1**), so it must be recognized that we were likely activating multiple presynaptic inputs and triggering recurrent, polysynaptic inputs. Third, our findings cannot directly inform us about whether these codes are actually used for computation *in vivo*. That requires behavioral responses from animals to determine whether downstream circuits utilize the information encoded with rate or synchrony^31^. Nonetheless, we believe that our findings are informative and can help to guide future work that attempts to examine potential divisions in coding in the neocortical circuit *in vivo*.

In summary, we found evidence that different subtypes of neurons in the neocortical microcircuit are differentially sensitive to information encoded with the synchrony or rate of presynaptic inputs. This suggests that the brain does indeed use both rate and timing codes, and may do so using different mechanisms and for different purposes.

## Materials and Methods

### Animals

Gin-GFP (FVB-Tg(GadGFP)45704Swn/J; JAX#003718) and Gad-GFP animals (CB6-Tg(Gad1-EGFP)G42Zjh/J; JAX#007677) were obtained from Jackson Laboratory. Mice were weaned at 21 days, in a temperature controlled room with a 12 hour light/dark cycle. Mice were given food and water, *ad libitum*. All procedures were in accordance with the regulations provided by the Canadian Council for Animal Care and approved by the Local Animal Care Committee at the University of Toronto Scarborough.

### Viral Microinfusion

Mice 5-7 weeks old received bilateral microinfusion of AAV1-CamKii-hChR2(H134R)-mCherry.WPRE.hGH (Addgene, #26975-AAV1) into layer 2/3 of the barrel cortex (−1.3mmAP, ± 3.1 mm ML, −1.1 mm DV). Mice were treated with ketoprofen (5 mg/kg) and anesthetized with isofluorane (4% induction, 2% maintenance). The anesthetized animal was then placed on a stereotaxic frame (Stoelting) and holes drilled in the skull above the coordinates of interest. To inject the viral vectors, a Hamilton Neuros Syringe (Hamilton, #65460-05) was connected to a microinjector (QSI, Stoelting) to infuse the virus at a volume of 0.15 uL per side with a rate of 0.05 uL/min. After each injection, the syringe was left in the brain for another 5 minutes to allow for sufficient diffusion of the virus. Following surgery, mice were treated with 0.5 ml of 0.9% saline subcutaneously and received ketoprofen post-operatively for 2-3 days.

### Immunohistochemistry

To confirm whether Gin-GFP and Gad67-GFP targeted SST+ and PV+ cells, respectively, brains from each transgenic line were fixed with 4% paraformaldehyde (PFA) via transcardial perfusion. After two days of fixation, the brains were sliced at 50 μm thickness using a vibratome (Leica).

For PV staining, free-floating brain sections of the barrel cortex were first washed in PBS and then incubated in 1% H_2_O_2_ in PBS for 30 minutes at room temperature. Slices were then blocked with PBS containing 10% goat serum, 3% bovine serum albumin, and 0.05% Triton-X-100 for 2 hours at room temperature. Afterwards, sections were incubated in PBS blocking buffer containing mouse anti-PV primary antibody (Thermofisher, 1:500) overnight at 4 °C. The next day, slices were washed in PBS and then incubated in PBS blocking buffer containing goat anti-rabbit secondary antibody conjugated with an Alexafluor 594 (Life Technologies, 1:500) for 1 hour at room temperature.

For SST staining, free-floating brain sections of the barrel cortex were first washed in PBS and then incubated in 1% H_2_O_2_ in PBS for 30 minutes at room temperature. Slices were then blocked with the same blocking solution as above for 2 hours at room temperature. Afterwards, sections were incubated in PBS blocking buffer containing mouse anti-SST primary antibody (Novus, 1:500) overnight at 4 °C. The next day, slices were washed in PBS and then incubated in PBS blocking buffer containing goat anti-rabbit secondary antibody conjugated with an Alexaflour 594 (Life Technologies, 1:500) for 1 hour at room temperature. Following this, tyramide signal amplification (TSA) was performed by incubating the sections in Rhodamine TSA reagent (1:30,000, diluted in 0.1 M Borate Buffer with 0.01% H_2_O_2_) for 30 minutes at room temperature.

Following staining, slices were washed with PBS, mounted onto gelatin-coated slides, and covered with a coverslip using Fluoshield (Sigma-Aldrich). Images were obtained using a confocal laser scanning microscope (Zeiss) with a 10x objective.

For cell counting experiments, L2/3 of the barrel cortex was imaged and was counted for GFP+, PV+ and SST+ cells. Approximately 4-6 sections/mouse were counted and averaged, with 4-6 mice/group. Genotypic specificity (total numbers of PV+ or SST+ cells / total numbers of GFP+ cells × 100), and efficiency (total numbers of GFP+ cells / total numbers of PV+ or SST+ cells × 100) were calculated using these averages.

### *Ex vivo* slice electrophysiology

Mice aged 7-12 weeks were anesthetized with 1.25% tribromoethanol (Avertin) and underwent cardiac perfusion using a chilled cutting solution containing (in mM): 60 sucrose, 25 NaHCO_3_, 1.25 NaH_2_PO_4_, 2.5 KCL, 0.5 CaCl_2_, 2 MgCl_2_, 20 D-glucose, 3 Na-pyruvate and 1 ascorbic acid, injected at a rate of approximately 1 mL/min. After 5-8 minutes of perfusion, the brain was quickly removed and cut coronally (350 μm thickness) with a vibratome (VT1000S) in chilled cutting solution to obtain slices of the barrel cortex. Once cut, these slices were then transferred into a recovery chamber comprising of a 50:50 mix of warm (34 °C) cutting solution and artificial cerebrospinal fluid (aCSF) containing (in mM): 125 NaCl, 25 NaHCO_3_, 1.25 NaH_2_PO4, 2.5 KCl, 1.3 CaCl_2_, 1MgCl_2_, 20 D-glucose, 3 Na-pyruvate, and 1 ascorbic acid. Following 30 min-1 hr of incubation, the slices were then transferred into an incubation chamber with room temperature aCSF. Within the recording chamber, aCSF was heated to 32 ^o^C using an in-line heater. Whole-cell current-clamp recordings were made using glass pipettes filled with (in mM): 126 K D-Gluconate, 5 KCl, 10 HEPES, 4 MgATP, 0.3 NaGTP, 10 Na-phosphocreatine. Glass capillary pipettes were pulled with a Flaming/Brown pipette puller with tip resistances between 4-8 MΩ.

### Patterned illumination with a digital multimirror device

To optically encode a 1-bit random signal into the activity of L2/3 pyramidal neurons, we used one-photon patterned illumination with a digital multimirror device (Polygon400, Mightex). After viral infusion surgery, and 2-3 weeks for expression (see above) we prepared slices and excited ROIs containing mCherry+ neurons in the slice. To target optical activation to as few neurons as possible, while maintaining reliable spiking responses in activated cells, we used software to draw circular patterns of 15 μm in diameter (PolyScan V2, Mightex) around mCherry+ cells. These patterns defined discs of illumination. The light used to activate the cells was generated from an LED with a 470 nm wavelength, and the optical power at the microscope stage was set to ∼14 mW/mm^2^, which we found to be sufficient for driving reliable spiking responses in targeted neurons.

To test the spatial specificity of our patterned illumination setup and determine whether it could reliably induce spiking, we conducted whole-cell patch clamp recordings from ChR2+ layer 2/3 pyramidal cells in the barrel cortex while illuminating circular patterns of blue light placed 25 μm apart in sequential order (see **Fig. S1**) in both the dorsal-ventral axis as well as the medial-lateral axis of the slice (470 nm, ∼14 mW/mm^2^). To determine the probability of spiking based upon the spatial distance of the light spot, the median probability of a spike occurring was calculated as well as the 95% confidence interval for the median via bootstrapping (*n* = 1000).

### Identifying ROIs with monosynaptic connections to a patched neuron

A criterion for including cell responses to rate and temporally coded signals was passing a test for reliable monosynaptic connections. Evidence for a monosynaptic connection was defined as a positive deflection in membrane potential 0.5-2.0 ms after the onset of optogenetic stimulation. This was characterized by fitting a piecewise linear function to the voltage with the form:

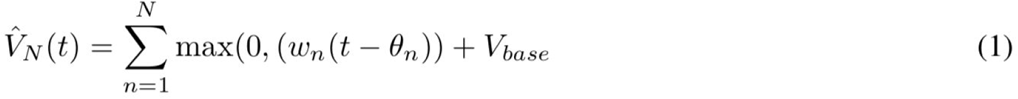

corresponding to the sum of N linear functions with slopes *w*_*n*_, with N different latencies *θ*_*n*_, and with a constant baseline V_base_. Functions with order 0 ≤ N ≤ 6 were fit. Functions of order N = 0 correspond to a constant baseline, i.e., no monosynaptic connection and no drift. Likewise, functions of order N = 1 correspond to a drifting baseline, i.e., no monosynaptic connection but a steady drift in voltage. Functions of higher order indicate a voltage deflection, i.e. a monosynaptic connection. First, the N = 0 and N = 1 models were compared in order to determine our null model of order N_null_. Specifically, a hypothesis test was performed to determine whether a N = 0 model should be rejected and a N = 1 model used as the null model for comparing to higher order (N ≥ 2) models. This hypothesis test was done with a chi-squared goodness of fit test:

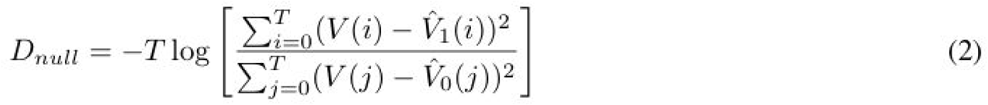

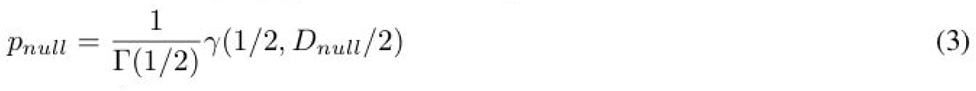

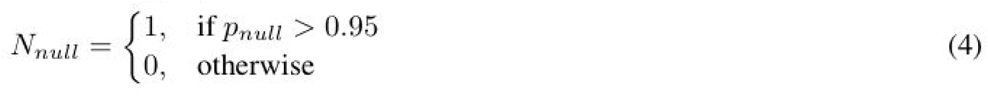

where Γ(*x*) corresponds to the gamma function, and γ(*x,y*) corresponds to the lower incomplete gamma function. In other words, if the likelihood of the data given N = 0 was less than 0.95, we rejected N_null_ = 0, and used N_null_ = 1. Again, to clarify, functions of order N ≥ 2 correspond to models with a monosynaptic connection. The model with the lowest Bayesian information criterion is our alternative model with order N_alt_ i.e.,

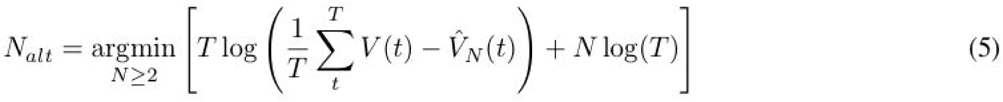

Subsequently, the alternative model 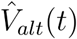 of order N_alt_ and the null model 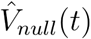 of order N_null_ are compared with a chi-squared goodness of fit test:

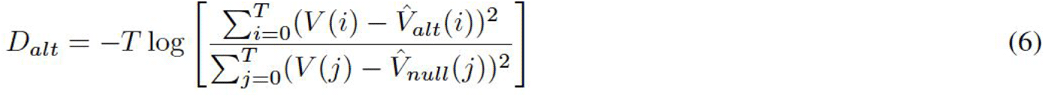

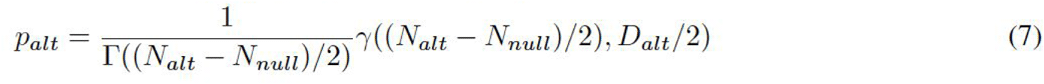

if *p*_*alt*_ ≥ 0.99, this was taken as evidence for a monosynaptic connection in a given trial.

Each of the 10 optogenetically targeted inputs underwent 5 trials for a total of 50 trials. From this, we calculated the number of trials where monosynaptic inputs were detected (i.e. where *p*_*alt*_ ≥ 0.99). If at least 90% of trials (45/50) showed a monosynaptic input according to this test (**Fig. 3C**) the cell was included in subsequent analyses. This allows for a degree of flexibility in the reliability of inputs, e.g., half of the optogenetically stimulated inputs could fail one trial, or one input could fail all trials.

### Encoding a 1-bit random signal with synchrony or rate of ROI activation

To examine the responses of different neuron types to synchrony and rate of inputs we developed protocols for optically encoding a random 1-bit signal (0 versus 1) in a brain slice. To do this, we drew 10 discs of illumination centred on mCherry+ neurons, that were in close proximity to the mCherry-patched cell. In order to mitigate unintended cross-stimulation of ROIs, we tried to space out the spots from each other by at least 50 μm, as we had previously observed that the probability of spiking dropped significantly if a spot was placed at least 50 μm distance from a cell (**Fig. S1**). A one-bit signal, s(*t*), was encoded in the optogenetically-driven activity of 10 presynaptic L2/3 pyramidal cells using either a rate or temporal code. Under a rate code, presynaptic neurons were driven by pulses 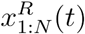 sampled from N independent inhomogenous Poisson process with rates, *λ*^*R*^(*t*), depending on the value of s(*t*), such that:

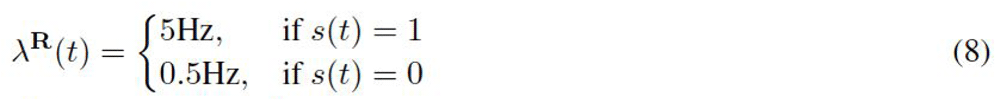

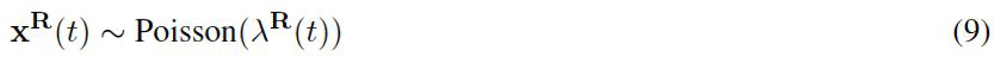

Under a synchrony code, presynaptic neurons were driven by pulses 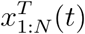 sampled from N independent homogeneous Poisson processes or from a single homogeneous Poisson process, creating states of uncorrelated and perfectly correlated pulses:

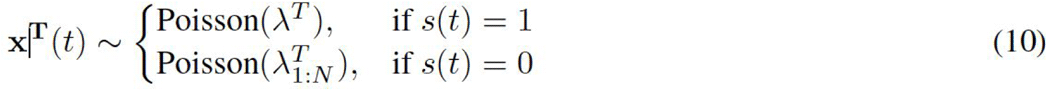

The rate *λ*^*T*^ was set to the mean rate for the rate-coded signal, i.e., *λ*^*T*^ *=* ⟨ *λ*^*R*^(*t*)⟩.

### Electrophysiological characterization

Electrophysiological characteristics for each neuron were estimated from 500 ms current injection steps (I_inj_) ranging from −80 pA to 400 pA in 40 pA increments. Eleven features were extracted in total including: (1) resting membrane potential (V_rest_, mV), (2) input resistance (R_in_, MΩ), (3) cell capacitance (C_mem_ pF), (4) membrane time-constant (τ_mem_, ms), (5) rheobase (I_θ_, nA), (6) f-I slope (f′, Hz/nA), (7) spike adaptation ratio, (8) sag amplitude (V_sag_, mV), (9) spike threshold (V_θ_, mV), (10) spike amplitude V_amp_, mV), and (11) spike half-width (T_half_, ms).

Standard calculations were used for these features^18,32,33^, but briefly, we will note the following for clarity:

- Spike times were identified as times at which membrane potentials crossed −20 mV with a positive gradient.
- Rheobase I_θ_ (current at which non-zero spike counts occur) and f-I slope, f’, were estimated by fitting the non-linear function *f*(*I*) = max(0, *f*′(*I* + *I*_θ_)to spike counts at each I_inj_ step value. Many cell types display spike accommodation with non-linear above-rheobase f-I relationships. We defined f-I slope to mean the initial slope above rheobase. Therefore, we fit this function to sub-rheobase and up to the first five above-rheobase spike counts inclusively.
- Spike adaptation ratio was estimated as the ratio between the last and first spike-time intervals (the difference between spike times). This requires a minimum of three spikes to estimate. For spike-trains with ≥ 7 spikes, the last two and first two intervals were used to estimate the ratio to improve estimate quality. Only the spike train from the highest I_inj_ was used to estimate this feature.
- V_rest_ was estimated as the average membrane potential in the 10 ms prior to current
- injection.
- V_sag_ was estimated as *V*_*sag*_ = | min(*V*) − *V*_*rest*_| at I_inj_= −80 nA during current injection.
- V_θ_ was estimated from all extracted spikes. A window around each identified spike time was used to extract action potential V(*t*) traces and the z-scored slope z(V’(*t*)) of each action potential was calculated. V_θ_ was estimated as the membrane potential at which z(V’(*t*)) ≥ 0.5
- V_amp_ was estimated as *V*_*amp*_ = | max(*V*) − *V*_*θ*_| for each action potential.
- T_half_ was defined as the duration of an action potential *V*(*t*) ≥ 0.5(*V*_*amp*_ −V_*θ*_) and was averaged across all extracted action potentials.
- R_in_ measurements were calculated and averaged across membrane potentials resulting from sub-rheobase, non-zero current injection. R_in_ was calculated using Ohm’s law as (*V*_*∞*_-*V*_*rest*_)/*I*_*inj*_ where the steady-state membrane potential V_∞_ was estimated as the mean membrane potential in the last 10 ms of current injection.
- τ_mem_ measurements were taken from membrane potential decay 100 ms after sub-rheobase non-zero current injection. Estimates were calculated by fitting a single order exponential function of the form *v* = (*v*_0_ − *u*) exp(−*t*/*τ*_*mem*_) + *u* and averaged.
- C_mem_ was calculated as τ_mem_/R_in_.

### Principal Components Analysis and Hierarchical Clustering

Principal components analysis of each cell’s electrophysiological characteristics was conducted using the sklearn python package^34^. To generate the dendrogram, we used the scipy.cluster package to implement hierarchical clustering^35^. Ward’s method was used to calculate the distance between each cluster.

### Mutual Information

Mutual information *I*(r;s)^36^ between the response *r(t)* of the postsynaptic neuron and the one-bit signal *s(t)* was used to assess the sensitivity of the postsynaptic neurons to the synchrony encoding of the signal:

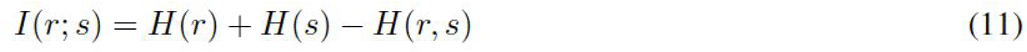

where *H*(r), *H*(s), *H*(r, s) are the entropy of the response, signal, and joint entropy of the signal and response respectively. The responses of the neurons, *r(t)*, were defined as mean membrane potential (e.g. **Fig. 5A & B**) or the spike counts (e.g. **Fig. 5C & D**) over the given temporal window.

Since temporal correlations increase amongst inputs when rates increase (**Fig. S4**), we could not directly calculate the mutual information between the rate of activations and the responses. Instead, the mutual information was conditioned on the number of presynaptic inputs 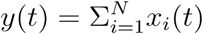 to discern how sensitive the postsynaptic neuron was to the rate-coded signal:

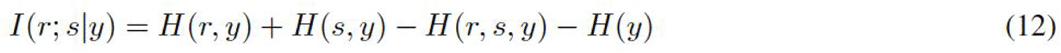

where H(r, y), H(s, y), and H(r, s, y) are the joint entropy of the postsynaptic response and number of presynaptic inputs, the signal state and the number of presynaptic inputs, and the response, signal state, and number of presynaptic inputs, respectively, whereas H(y) was the entropy of the number of presynaptic inputs.

To estimate the entropy of discrete variables^36,37^ U (such as signal state, number of presynaptic input, postsynaptic spike count) we used:

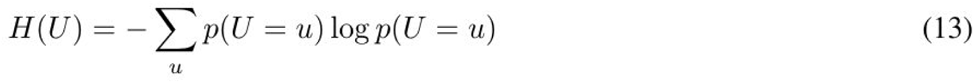

The entropy of continuous variables (average membrane potential) was estimated by constructing histograms to approximate the probability density function of the variable, i.e.:

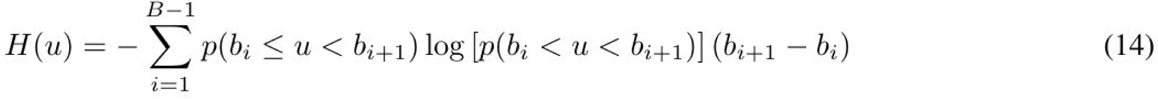

Along bin edges b, where the number of bins (*N*_*bins*_ *= B-1*) was chosen as the maximum of Sturges’ formula^38^ and the Freedman-Diaconis rule^39^:

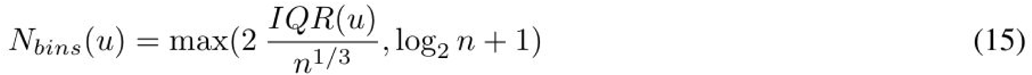

Where *n* is the size of data *u*.

## Data Analysis and Statistics

All data analysis code was written in python 2.7 using tools from the scientific computing ecosystem (numpy^32^, scipy^33^, matplotlib^34^, scikit-learn^35^, pandas^36^). <u>All code and all data will be made publicly available upon acceptance for publication.</u>

To determine whether the mean mutual information was different between each cell type, we ran a One-Way ANOVA, or Kruskal-Wallis test if Levene’s test indicated unequal variances between groups. We also applied post-hoc individual t-tests for differences between pairs of groups, or Welch’s t-test if Levene’s test indicated unequal variance. Bonferroni corrections were applied for each test. All tests were run using scipy.

## Contributions

BAR, MMK and JK conceived of the experiments and analysis. MMT, DG, LS and HC collected the data. MMT and LYP analyzed the data. MMT, LYP and BAR wrote the manuscript. MMT, LYP, MMK, JK and BAR edited the manuscript.

## Acknowledgements

This work was supported by a Human Frontier Science Program Young Investigator Grant to MMK, JK and BAR (RGY0073/2015) and a Natural Sciences and Engineering Research Council of Canada Discovery Grant to BAR (RGPIN-2014-04947).

## Supplemental Figures

**Supplemental Figure 1.**
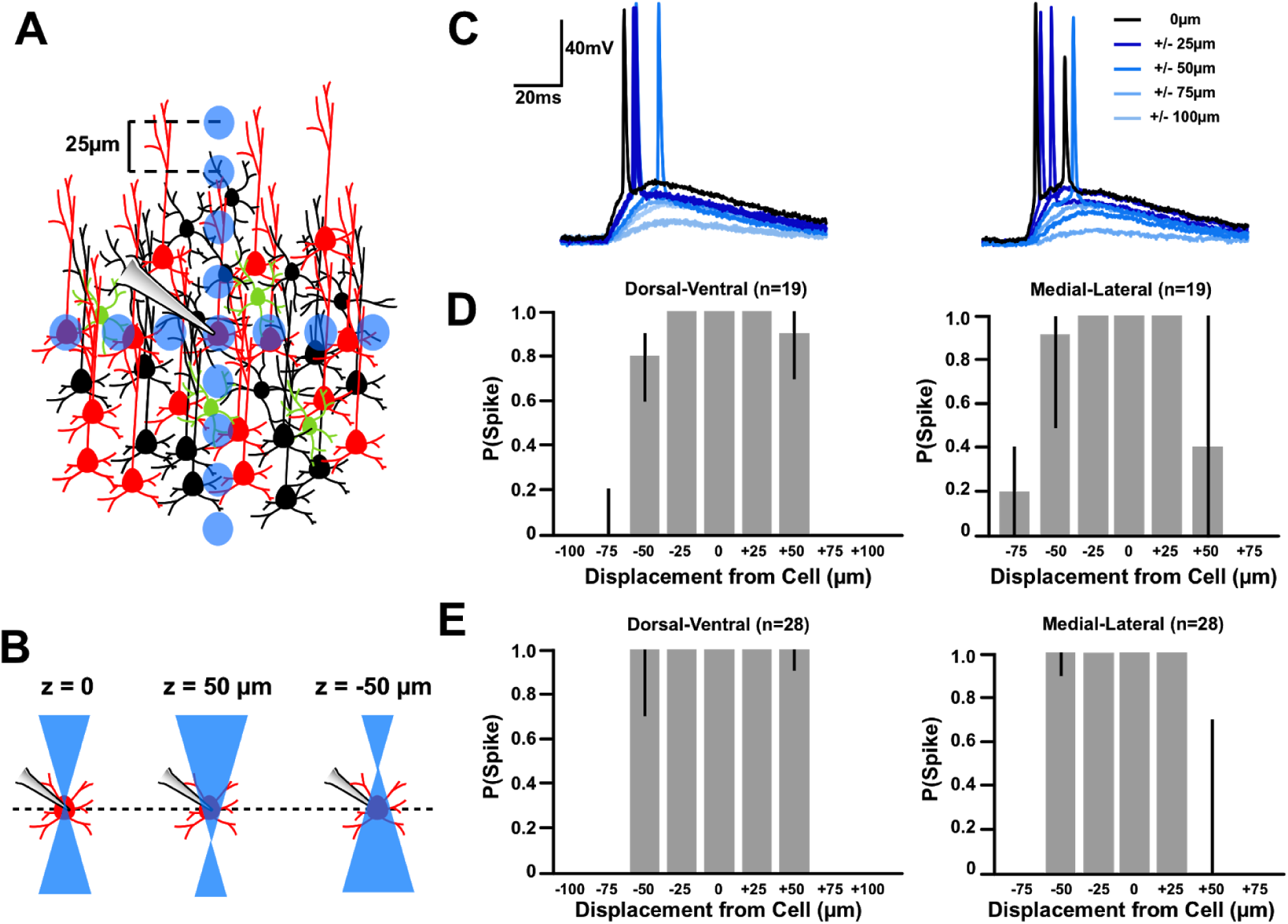
L2/3 Pyramidal Neurons reliably spike when illuminated with 15 μm spots placed ≤50 μm away. A) Experimental procedure. B) Given that single-photon illumination produces a cone of illumination above and below cells being stimulated, we conducted the protocol shown in A), while also stimulating the recorded cell when the light was in focus and directly on top of the cell (z = 0), above it (z = 50 μm) or below it (z = −50 μm). C) Sample responses from ChR2+ neuron to spots placed at varying locations when the light was directly focused on the cell (z = 0) D) Median spiking probability of ChR2+ neurons (n= 15) to spots placed at varying locations within the microscope’s field of view with spot directly in focus of the cell (i.e. z = 0). E) Same analysis as D) except with cellular responses to spots illuminated when the focal point was above or below the cell of interest (i.e. z = 50 μm or z = −50 μm).

**Supplemental Figure 2.**
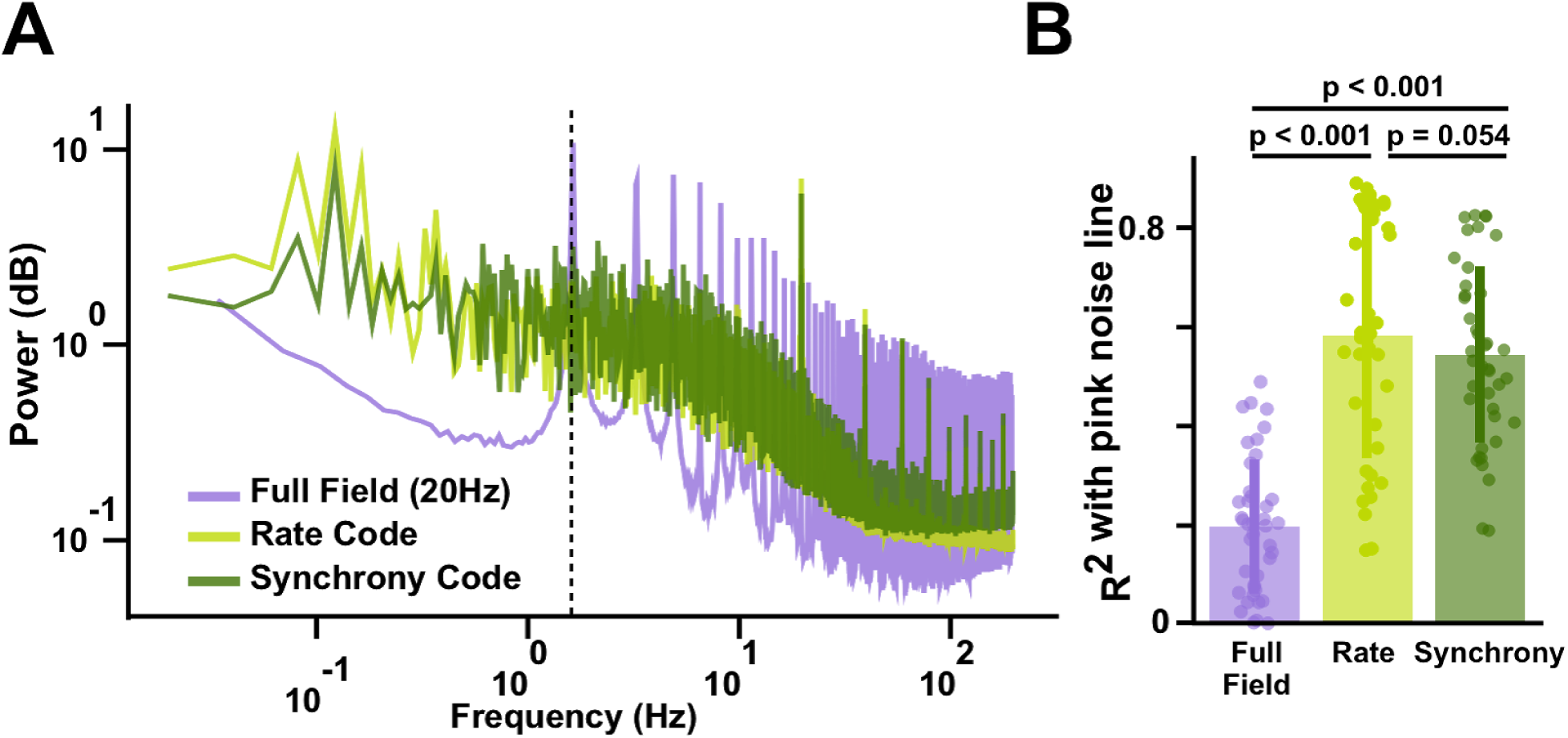
Artificial rate and synchrony of optical inputs to Layer 2/3 pyramidal neurons mimic *in vivo*-like responses in postsynaptic cells. A) Power spectral density (PSD) of neuronal recordings to either full field, rate or synchrony of optical inputs. B) Correlations of each spectrum to a pink noise line.

**Supplemental Figure 3.**
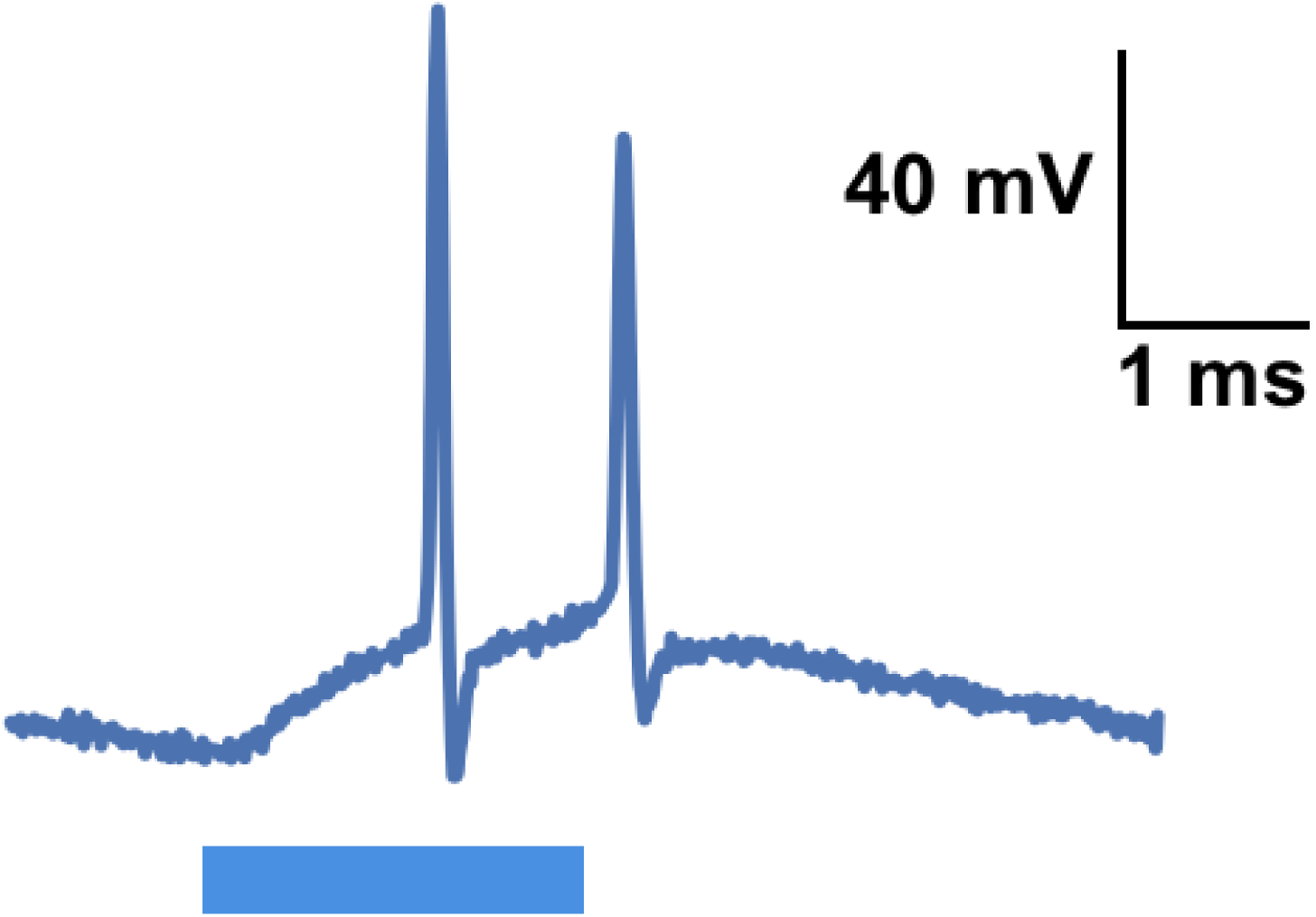
Prolonged polysynaptic activity was produced upon activation of our ROIs.

**Supplemental Figure 4.**
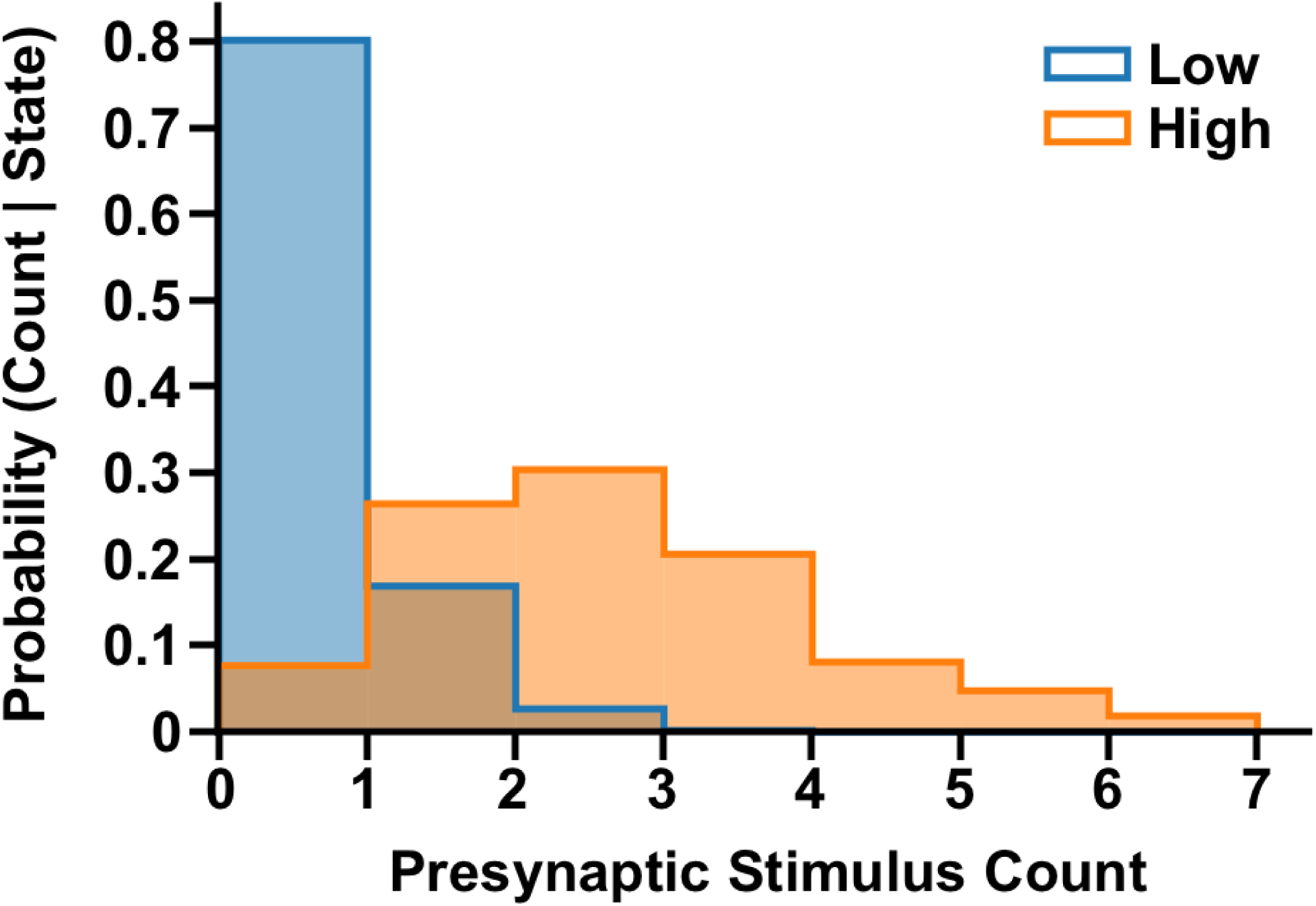
Presynaptic spike count increased when higher rates of optical input were used.

